# Connecting coil-to-globule transitions to full phase diagrams for intrinsically disordered proteins

**DOI:** 10.1101/2020.05.13.093351

**Authors:** X. Zeng, A. S. Holehouse, T. Mittag, A. Chilkoti, R. V. Pappu

## Abstract

Phase separation is thought to underlie spatial and temporal organization that is required for controlling biochemical reactions in cells. Multivalence of interaction motifs also known as stickers is a defining feature of proteins that drive phase separation. Intrinsically disordered proteins with stickers uniformly distributed along the linear sequence can serve as scaffold molecules that drive phase separation. The sequence-intrinsic contributions of disordered proteins to phase separation can be discerned by computing or measuring sequence-specific phase diagrams. These help to delineate the combinations of protein concentration and a suitable control parameter such as temperature that support phase separation. Here, we present an approach that combines detailed simulations with a numerical adaptation of an analytical Gaussian cluster theory to enable the calculation of sequence-specific phase diagrams. Our approach leverages the known equivalence between the driving forces for single chain collapse in dilute solutions and the driving forces for phase separation in concentrated solutions. We demonstrate the application of the theory-aided computations through calculation of phase diagrams for a set of archetypal intrinsically disordered low complexity domains.

**STATEMENT OF SIGNIFICANCE:** Intrinsically disordered proteins that have the requisite valence of adhesive linear motifs can drive phase separation and give rise to membraneless biomolecular condensates. Knowledge of how phase diagrams vary with amino acid sequence and changes to solution conditions is essential for understanding how proteins contribute to condensate assembly and dissolution. In this work, we introduce a new two-pronged computational approach to predict sequence-specific phase diagrams. This approach starts by extracting key parameters from simulations of single-chain coil-to-globule transitions. We use these parameters in our numerical implementation of the Gaussian cluster theory (GCT) for polymer solutions to construct sequences-specific phase diagrams. The method is efficient and demonstrably accurate and should pave the way for high-throughput assessments of phase behavior.

## INTRODUCTION

Intrinsically disordered proteins that have the requisite valence of adhesive linear motifs – stickers – can drive phase separation and give rise to membraneless biomolecular condensates [1-5]. Knowledge of how phase diagrams vary with amino acid sequence and changes to solution conditions is an essential component of understanding how proteins contribute to condensate assembly and their regulated dissolution [1-4, 6]. Comparative assessments of sequence-to-phase diagram relationships are also invaluable for enabling the design of novel disordered low complexity domains (LCDs) that give rise to synthetic condensates for engineering applications [7-11]. In this work, we introduce a two-pronged computational approach to predict sequence-specific phase diagrams. We start by extracting key parameters from simulations of single-chain coil-to-globule transitions. We use these in our numerical implementation of the Gaussian cluster theory (GCT) for polymer solutions [12, 13] to construct sequences-specific phase diagrams.

Our approach leverages the known equivalence between the driving forces for coil-to-globule transitions of individual polymer molecules in dilute solutions and phase separation that arises from collective interactions among polymer molecules in concentrated solutions [10, 12-17]. For systems with an upper critical solution temperature (UCST) (**Figure 1**), the coil-to-globule transition is characterized by indifferent solvent conditions at the theta temperature (*T*_θ_), poor solvent conditions below *T*_θ_, and good solvent conditions above *T*_θ_ [11]. At finite concentrations, for *T* < *T*_θ_, chains overlap with one another, and because of cohesive interactions among chain molecules there exist temperature-dependent threshold volume fractions above which the system separates into two phases namely, a dilute solution that coexists with a dense phase. For a temperature *T* < *T*_θ_, we denote the volume fractions of the coexisting dilute and dense phases as ϕ_b1_ and ϕ_b2_, respectively (**Figure 1**). While the dilute arm of the binodal namely ϕ_b1_ as a function of *T* can be obtained via a range of experimentally tractable approaches, accurate estimates of the dense phase concentration *i.e*., ϕ_b2_ as a function of *T* are plagued with several challenges. These include the need for high concentrations of protein, artefacts from fluorescent reporters, the impact of finite size effects, and slow equilibration within dense phases [7]. Accordingly, despite their over-arching importance for characterizing condensates, measurements of system-specific binodals are available for only a handful of LCDs [3, 8-10, 18].

**Figure 1:**
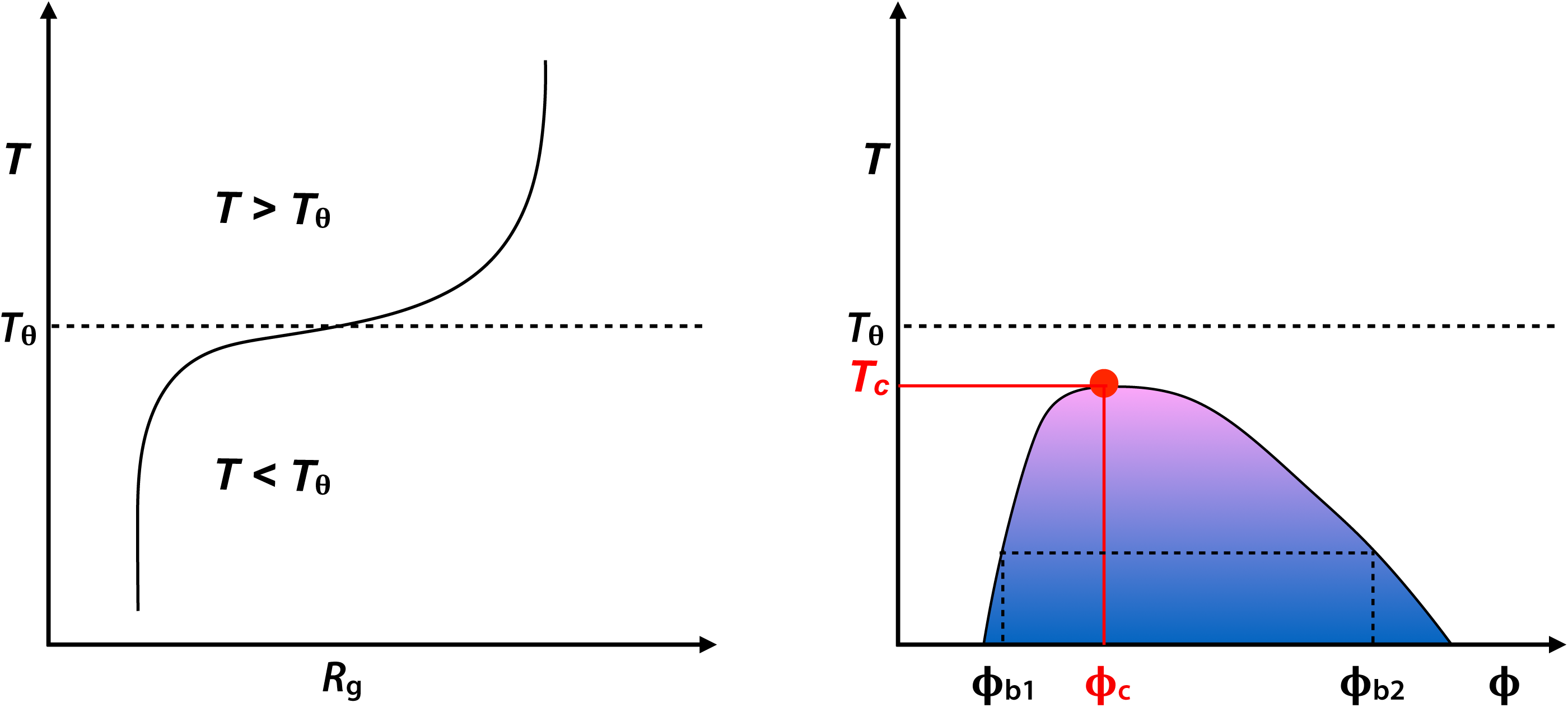
Equivalence between single chain coil-to-globule transitions and phase separation. The schematic depicts expectations for a system that shows UCST behavior. (A) Coil-to-globule transitions whereby coils form at temperatures above *T*_θ_ (dashed line) and globules form below *T*_θ_. At *T*_θ_ the radius of gyration scales as *N*^0.5^ with chain length *N*. There is a system-specific collapse temperature *T*_collapse_ below which *R*_g_ ∼ *N*^0.33^. The crossover regime, whose width is defined by the ratio of the two- and three-body interaction coefficient, corresponds to the interval *T*_collapse_ < *T* < *T*_θ_ where the scaling exponent for *R*_g_ lies between 0.33 and 0.5. (B) Expected phase diagram for a flexible polymer that shows UCST behavior. The critical point defined by the critical temperature (*T*_c_) and critical volume fraction (ϕ_c_), is the point where the binodal and spinodal coincide. The binodal is shown as the contour that delineates the two-phase regime (shown in colored shading, where the colors become cooler away from the critical point). For a given temperature *T* < *T*_θ_ and *T* < *T*_c_, there exists a volume fraction above which the system separates into two coexisting phases, a dilute and dense phase characterized by volume fractions ϕ_b1_(*T*) and ϕ_b2_(*T*). The difference between ϕ_b1_(*T*) and ϕ_b2_(*T*) decreases as *T* approaches *T*_c_.

The driving forces for temperature-dependent coil-to-globule transitions and phase separation are governed by the three-way interplay among chain-chain, chain-solvent, and solvent-solvent interactions [14]. The consequences of this three-way interplay are captured via an effective two-body interaction coefficient *B*, which has units of inverse volume. The value of *B* is positive below *T*_θ_ for a system with UCST behavior and negative for a system above *T*_θ_ for a system with a lower critical solution temperature (LCST). These features of *B* signify net attractions among monomer units below *T*_θ_ for a UCST system and above *T*_θ_ for a LCST system. The three-body interaction coefficient *w* (**Figure 2**) is always positive and to first approximation it is independent of temperature [14]; the magnitude of *w* is determined, in part, by the local persistence length or thickness of the chain. The steepness of coil-to-globule transitions and the temperature-dependent values of ϕ_dense_ are governed by the ratio of the two- to three-body coefficient [12, 13].

**Figure 2:**
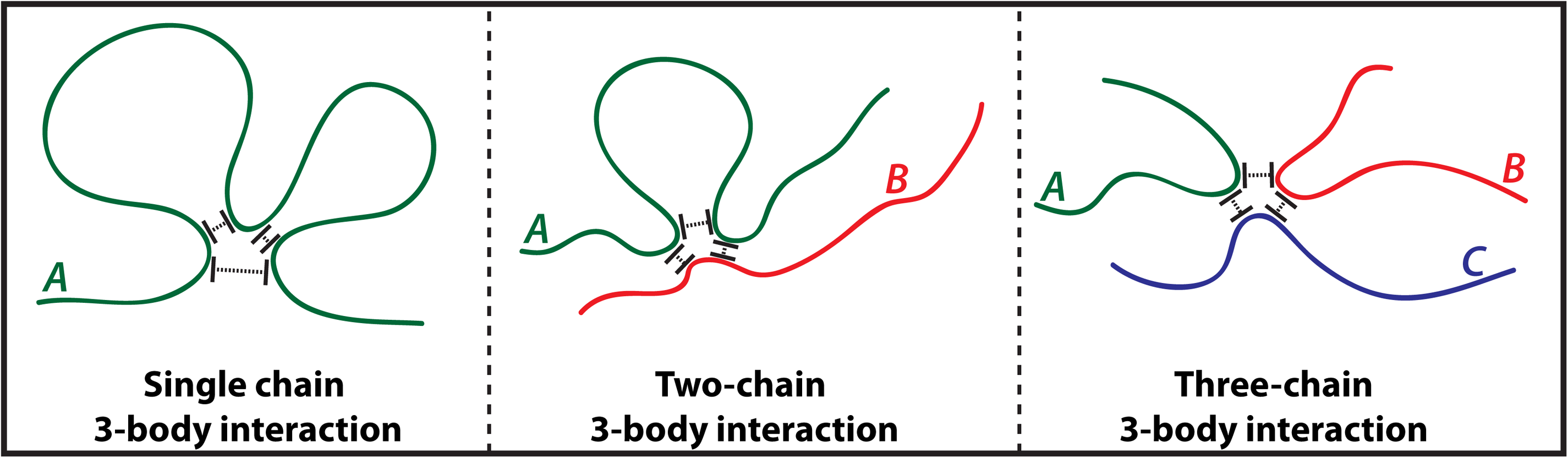
Depicting how three-body interactions are realized. Three body interactions are realizable in at least three different ways: within a single polymer (left), between pairs of polymers (middle), among three polymers (right). The interactions are always repulsive, and this derives from the finite thickness and steric impenetrability of polymers.

Disordered LCDs (as well as folded domains) can be mapped onto *sticker*-and-*spacers* architectures [10, 19-23]. Stickers are attractive groups [24, 25] that include short linear motifs or individual residues that form reversible physical (non-covalent) crosslinks with one another; conversely, the excluded volumes of spacers control the extent of compaction of individual chains and the cooperativity of phase transitions that give rise to two coexisting phases [23]. Recent studies have shown that the correspondence between coil-to-globule transitions and phase separation, which is well established for flexible homopolymers, is also applicable for describing the phase behavior of disordered LCDs [16, 26]. In order for this correspondence to apply, the stickers have to be uniformly distributed along the linear sequence [10]. Intriguingly, a significant fraction of naturally occurring disordered LCDs, specifically prion-like domains, and designed sequences have the requisite sequence features that should enable correspondence between the driving forces that control coil-to-globule transitions and phase separation [9-11, 27-29].

We reasoned that it should be possible to compute complete phase diagrams using quantitative analysis of sequence-specific coil-to-globule transitions in conjunction with a requisite theoretical framework that directly connects coil-to-globule transitions to phase separation. Accordingly, we deployed a two-pronged strategy for calculating sequence-specific phase diagrams of disordered LCDs. First, all-atom or coarse-grained simulations are used to obtain temperature-dependent coil-to-globule transitions and the results are analyzed to extract *T*_θ_, *B*, and *w*. Next, these parameters are used in our numerical adaptation of the GCT for polymer solutions [12, 13] in order to construct full phase diagrams in the two parameter (ϕ,*T*) space.

The GCT was developed for generic homopolymers, each comprising *N* monomers connected by *N*–1 bonds of length *l*_0_. The distributions of intra-molecular and inter-molecular distances are described in terms of distances between statistical Kuhn segments instead of monomers. At their specific theta temperatures, any homopolymer can be reduced to a freely jointed chain of Kuhn segments, where the latter refers to the length scale beyond which the chemical details of the repeating units can be ignored [30]. If *n*_s_ is the number of bonds per statistical Kuhn segment, then the number of Kuhn segments is defined as *n*_K_ = (*N*–1)/*n*_s_ and mean squared length of each Kuhn segment is written as 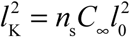 where *C*_∞_ is the characteristic ratio of the chain [30]. The GCT assumes Gaussian forms for the distributions of intra-chain and inter-chain distances between Kuhn segments [12, 13]. The main parameters of the Gaussian distributions are functions of *T, T*_θ_, *B*, and *w*. Mean-field free energies, written in terms of moments of the Gaussian distributions, describe the temperature-dependence of intra-chain and inter-chain interactions. Minimization of the sum of single- and multi-chain free energies allows us to extract temperature-dependent values of 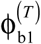 and 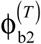. The locus of these temperature-dependent values defines the coexistence curve or *binodal*; likewise, the locus of points where the second derivative of the sum of single- and multi-chain free energies is zero helps define the instability curve or *spinodal*. In this way, we can compute system-specific full phase diagrams by minimization of the overall free energy for a polymer solution based on parameters obtained from single chain coil-to-globule transitions.

The remainder of the manuscript is organized as follows: First, we summarize the main tenets of the GCT. Next, we detail how simulations of coil-to-globule transitions of individual LCDs are used to extract parameters for use in the GCT and demonstrate how we use our numerical implementation of the GCT to compute sequence-specific, temperature-dependent binodals, and spinodals. We then illustrate the accuracy of our two-pronged approach by computing phase diagrams for LCDs where temperature-dependent binodals have been measured. We further illustrate the methodology by computing full phase diagrams for distinct synthetic systems that have upper and lower critical solution temperatures, respectively. We conclude with a discussion of how the GCT-aided two-pronged method might be useful for high-throughput characterizations of phase diagrams of disordered LCDs.

## MATERIALS AND METHODS

### Systems of interest

We demonstrate the application of GCT-aided calculation of phase diagrams using simulations of coil-to-globule transitions for a series of systems. We start with the 137-residue LCD from the hnRNP-A1 protein [31]. Recently, Martin et al. [10] characterized the phase behavior of this system and three designed variants that differ from the wild-type sequence in terms of the number (valence) of aromatic stickers. The sequences of interest are referred to as WT-A1, Aro^+^ that was designed to have higher valence of aromatic residues than WT-A1, and Aro^-^ as well as Aro^--^ that have lower numbers of aromatic stickers than the WT [10]. For these systems, we use coarse-grained simulations of coil-to-globule transitions that are based on PIMMS [10, 32], which is a lattice-based simulation engine.

Polyglutamine (polyQ) expansions are associated with Huntington’s disease and at least eight other neurodegenerative disorders [33]. Considerable effort has focused on characterization the conformational equilibria of individual polyQ chains and the phase equilibria of these systems [17, 34-49]. Homopolymers of polyQ forms globules in aqueous solvents and experiments as well as simulations point to UCST phase behavior [34, 42, 49, 50]. Additionally, evidence has been presented for the prospect of disordered, oligomeric / liquid-like phases serving as precursors of fibril formation [34, 35, 49, 51, 52]. Here, we use the ABSINTH implicit solvation model [53] to perform all-atom simulations of coil-to-globule transitions. We use these simulations to demonstrate how the GCT can be used to construct phase diagrams for a biologically relevant system of homopolypeptides.

Systems that undergo thermoresponsive phase behavior are of considerable interest for the design of biomaterials [9, 27-29]. As a final set of systems, we demonstrate the applicability of the two-pronged approach for constructing temperature-dependent phase diagrams of two systems *viz*., (QGQSPYG)_9_ and (TPKAMAP)_9_. These synthetic systems are designed to have UCST and LCST behaviors, respectively.

### Coarse-grained and all atom simulations of coil-to-globule transitions

PIMMS is a lattice-based simulation engine that uses a single-bead per amino acid residue [10, 32]. Simulations of coil-to-globule transitions of WT-A1, Aro^+^, Aro^-^, and Aro^--^ were performed using a stickers-and-spacers representation of these sequences. Here, aromatic residues are stickers and all other residues are spacers. In units of *k*_*B*_*T* = 1 where *k*_*B*_ is Boltzmann’s constant, Martin et al. showed that sticker-sticker energies of –14, sticker-spacer energies of –5, and spacer-spacer energies of –2 provide a minimalist model that reproduces the variation of *R*_g_ with sticker valence in dilute solutions and experimentally measured binodals in the space of ϕ and *T*, with suitable conversions between the reduced temperature units and Kelvin [10]. In this work, we use results from PIMMS simulations as the touchstone for comparative assessments of GCT-derived phase diagrams and we use unitless, reduced temperatures, setting *k*_*B*_*T* = 1 for comparative calculations on the four systems, WT-A1, Aro^+^, Aro^-^, and Aro^--^.

The all atom simulations for polyQ sequences denoted as Q_n_ (n=20, 30, 40, 50, 60, 70) and the two sequences (QGQSPYG)_9_ and (TPKAMAP)_9_ were performed using version 2.0 of the CAMPARI modeling suite (http://campari.sourceforge.net/). The N- and C-termini are capped using N-acetyl and N′-methylamide groups, respectively. All simulations use the ABSINTH implicit solvent model and force field paradigm. The parameters were derived based on OPLS-AA/L forcefield and all the parameters are implemented in abs_3.2_opls.prm parameter set. Parameters for proline residues were based on the work of Radhakrishnan et al. [54]. The cutoff for the short-range Lennard-Jones potential is set to be 10 Å. The cutoff for electrostatic interactions between different sites on neutral groups is 14 Å. There are no cutoffs among charge groups that carry a net charge, and these pertain to solution ions as well as charge groups on the sidechains. In systems with charged residues, specifically (TPKAMAP)_9_, nine Cl^-^ ions, modeled using the parameters developed by Mao and Pappu [55], were included to ensure electroneutrality. In addition, ion pairs were included to mimic an excess NaCl concentration of 1 mM. For the temperature dependence of macroscopic dielectric constant of water we use the form adapted by Wuttke et al. [56].

ABSINTH uses experimentally derived reference free energies of solvation for model compounds [53, 57]. The references free energies of solvation have weak temperature dependencies for polar compounds. Accordingly, we use the reference values at 298 K for the model compounds that mimic sidechains of Gln, Ser, Gly, Pro, and Tyr. However, to model LCST behavior for (TPKAMAP)_9_, we adapt the generalization introduced by Wuttke et al. [56]. Specifically, we set the reference free energies of solvation for the backbone amide and the model compounds that mimic the sidechains to be temperature dependent. The parameters for the enthalpy and heat capacity that are used in the Gibbs-Helmholtz equation to estimate temperature dependent free energies of solvation are summarized in **Table S1**.

Standard Metropolis Monte Carlo simulations (MMC) aided by thermal replica exchange [58] were used to sample conformations of polyQ systems and the two synthetic sequences. Move sets for MMC include side chain torsions, concerted rotations, pivot moves, translations of ions, and translations and rotations of individual molecules [10, 54, 57, 59]. Details regarding the number of replicas and the temperature schedule for each system are listed in **Table S2**. Swaps between neighboring replicas were proposed once every 5×10^4^ MMC steps. Simulations were initiated using a randomly generated self-avoiding conformation for each polypeptide. All simulations use a spherical droplet and the droplet is large enough to ensure against confinement artefacts. Multiple independent replica exchange simulations were performed – three for (TPKAMAP)_9_ and four for the other systems – and each independent replica exchange simulation comprises a total of 6×10^7^ MMC steps.

## THEORY

### Gaussian cluster theory (GCT) for a single chain

The workflow for constructing full phase diagrams using simulations of single chain coil-to-globule transitions and the use of GCT based on parameters extracted from the single-chain simulations is summarized in **Figure 3**. We first connect the GCT formalism to results from single-chain simulations and show how this connection allows us to estimate *T*_θ_, the two-body interaction coefficient *B*, and the three-body interaction coefficient *w*. The GCT prescribes specific forms for the distribution of intra-chain and inter-chain distances between chain segments [12, 13]. The segments of interest are Kuhn segments. Formally, a real chain of *N* residues can be reduced to an effective freely jointed chain of *n*_K_ Kuhn segments. For polypeptides, the length of a Kuhn segment can be calculated using *l*_0_ =3.8Å as the length of a virtual bond between adjacent alpha carbon atoms for all-atom simulations and *l*_0_ =1 lattice unit for PIMMS simulations. We set *n*_K_ to be five for polyQ and the WT-A1 system as well as variants of this system. This choice is a robust estimate for systems that have the local flexibility of generic polypeptides, which refers to systems where proline residues make up less than 10% of the sequence [60]. For the synthetic systems, we set *n*_K_ to be 7, which is identical to the number of residues within each repeat. Systematic calibrations showed that the calculated phase diagrams are relatively insensitive to the choice made for *n*_K_.

**Figure 3:**
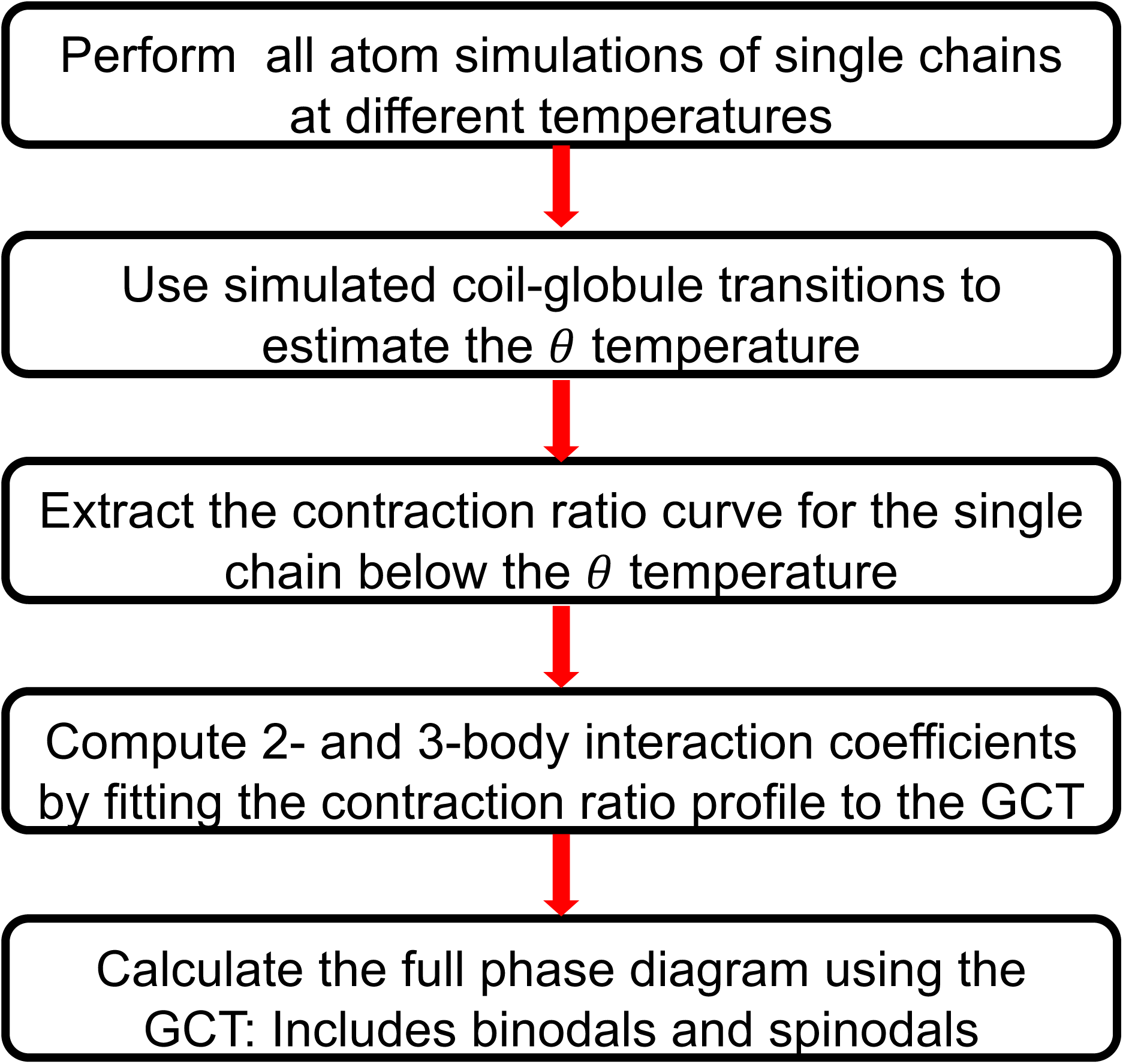
Workflow summarizing the overall two-pronged approach that leverages the GCT formalism. We use the prescribed approach to connect results from single chain simulations of coil-to-globule transitions to calculations of phase diagrams using the GCT.

The GCT assumes a Gaussian form for ρ(*r*), the distribution of inter-segment distances within a single chain [12, 13], which is written as:

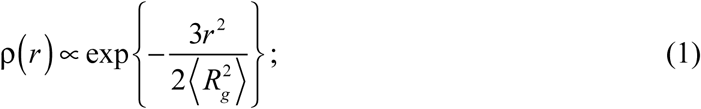

Here, 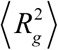 is the ensemble-averaged value of the mean-squared radius of gyration for a single chain. To simplify the algebra, the theory introduces three parameters *viz*., the reduced temperature τ, the parameter σ that quantifies the standard deviation of the Gaussian distribution for ρ(*r*), and the contraction ratio α_s_:

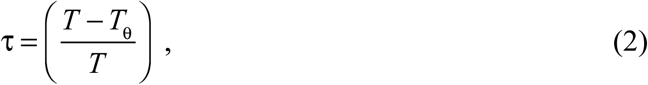

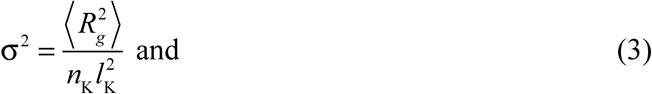

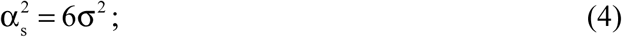

Notice that τ < 0 for *T* < *T*_θ_. Allegra and Ganazzoli [61] showed that the strain ratio for low wavenumbers (*k*) in Fourier space, which translates to large separations *r* in real space, can be defined as: 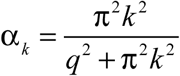. Accordingly, the strain ratio in Fourier space, and the contraction ratio in real space depend on a single auxiliary parameter *q*, known as the *compression parameter* [12, 13]. Here, *q* tends to zero as *T* approaches *T*_θ_ whereas *q* approaches ∞ in the non-physical limit where the chain is compressed down to a single point. Based on the relationship between ρ(*r*) and its Fourier transform, which is also a Gaussian distribution in *k-*space, we rewrite ρ(*r*) and *r* in terms of *q*; moments such as 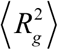 can be evaluated in terms of *q*. Allegra and coworkers have shown that [61, 62]:

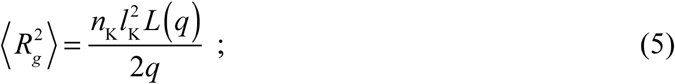

Here, *L*(*q*) is the Langevin function 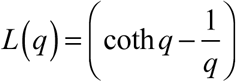. From equations (3) – (5) it follows that 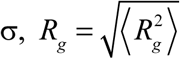 and the contraction ratio α_s_ are functions of *q*.

The mean-field free energy *A*_1_(*q*; τ) of a single chain at a relative temperature τ is written in terms of the compression parameter *q*. Given that *q* is a continuous parameter that quantifies the dependence of chain dimensions on *T*, we have an analytical route to relate the temperature-dependent chain dimensions to the free energy of the chain. At equilibrium, the free energy of a single chain is minimized by setting 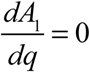. The desired free energy *A*_1_(*q*; τ) is a sum of the effects of intra-chain interactions *A*_intra_(*q*; τ), and the elastic term *A*_el_(*q*) that quantifies contributions of conformational entropy. Accordingly,

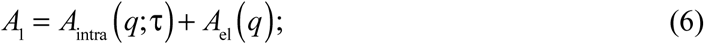

From the assumption of a Gaussian distribution for intra-segment distances within a chain, it follows that *A*_intra_(*q*; τ) is written in terms of *q*, τ, the two-body interaction coefficient *B* (where *B* > 0), and the three-body interaction coefficient *w*, as:

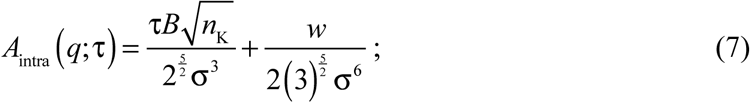

The parameter τ*B* is the second virial coefficient that reflects the interplay of pairwise segment-segment, segment-solvent, and solvent-solvent interactions. Note that for UCST systems, τ*B* is negative for *T* < *T*_θ_ because *B* here is positive; conversely, for LCST systems, *B* is negative and the second virial coefficient τ*B* is negative for *T* > *T*_θ_. The parameter *w* is a direct measure of the third virial coefficient defined by three-way interactions of polymer segments (**Figure 2**).

The elastic free energy is written in terms of the compression parameter *q* as:

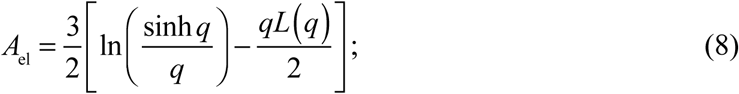

At equilibrium, the free energy of a single chain at fixed τis minimized by setting 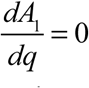 This yields a relationship between τ and the compression parameter *q*, which allows us to estimate *q* by solving the transcendental equation below:

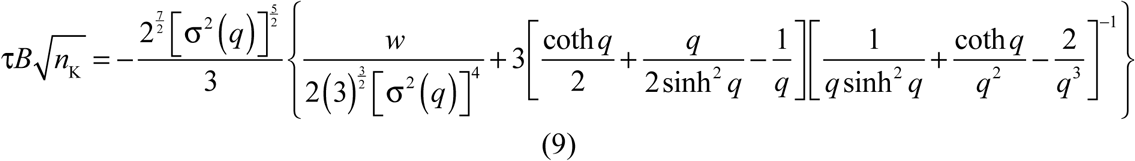

### Extraction of *T*_θ_, *B*, and *w* by comparing GCT to results from single-chain simulations

Equation (9) provides the central connection between simulations of coil-to-globule transitions and parameters of the GCT. The protocol for its usage is as follows:

1. Coarse-grained or all-atom simulations are used to extract the system-specific profile for 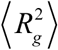 as a function of simulation temperature *T*. This information is converted into a profile for the contraction ratio α_s_ versus *T* using equations (3) and (4). Additionally, for a given temperature we extract internal scaling profiles, which plot ⟨ ⟨ *R*_ij_ ⟩ ⟩ versus |j–i|. Here, ⟨ ⟨ *R*_ij_ ⟩ ⟩ refers to ensemble-averaged mean distance between all pairs of residues i and j that are |j–i| apart along the linear sequence.
2. If the system of interest is a perfect homopolymer or can be reduced to a perfect homopolymer, then simulations of coil-to-globule transitions are performed for a series of chain lengths or numbers of repeats. In this scenario, *T*_θ_ is estimated as the intercept along the temperature axis where plots of 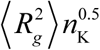 versus *T* intersect with one other. This is reflective of the tricritical nature of the theta point [12, 13, 63, 64]. Alternatively, given temperature-dependent profiles for plot ⟨ ⟨ *R*_ij_ ⟩ ⟩ versus |j–i|, we identify *T*_θ_ as the temperature for which ⟨ ⟨ *R*_ij_ ⟩ ⟩= *R*_0_ |j–i|^0.5^ [65].
3. Next, we fit a smooth curve to the profile of α_s_ versus *T*. From this profile, we estimate *T*^*\**^, defined as the temperature where 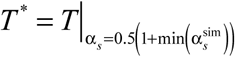. Here, min 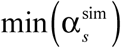 is the minimum value of the contraction ratio from the simulations. For the UCST system, min 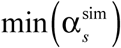 is the contraction ratio at the lowest sampled temperature. For the LCST system, it is the value of the contraction ratio at the highest sampled temperature.
4. From the GCT, it follows that the transition temperature has to satisfy 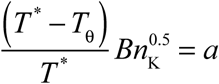 Accordingly, given prior knowledge of *T*_θ_ and *T*^*\**^ we can estimate the value of *B*. Raos and Allegra have shown that *a* ≈ –0.35 for generic homopolymers [12]. We use this to set bounds on the value of *a* and develop a numerical procedure (**Figure S1**) to estimate *T*^*^.
5. The relationship in equation (9) is used to rewrite the compression parameter *q* as a function of τ. This, in conjunction with the definitions introduced in equations (3) and (4), leads to numerical profiles for the contraction ratio as a function of 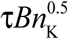.
6. We then estimate *w* using a parameter scan where the constraint is the requirement that the theoretical profile for the contraction ratio versus 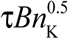 be coincident with the profile derived from the single-chain simulations.

In practice, we identify a putative transition region for the location of *T*^*\**^, choose different values of *T*^*^ in this region, and simultaneously choose values for *B* and *w*, using the bounds based on the constraint prescribed in step (4). We then identify the combination that minimizes the difference between the simulated contraction ratio and the one calculated using equation (9) from the GCT. We have developed two distinct approaches for estimating *B* and *w* and these approaches are described in the pseudo code shown in **Figure S1**. It helps to have roughly ten distinct temperatures that span the transition region and the region where the contraction ratio plateaus as the chain forms a globule.

### GCT for phase separation used to calculate full phase diagrams

The GCT assumes that in a cluster comprising *v* distinct polymers, the distribution of distances for polymer *i* vis-à-vis all other polymers is a Gaussian of the form [12, 13]:

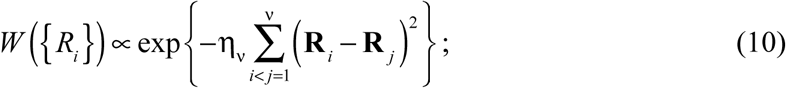

In equation (10), **R**_*i*_ and **R**_*j*_ denote the centers-of-mass of polymers *i* and *j*; the parameter 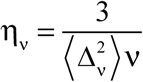 is related to the ensemble-averaged mean square distance between chains, 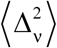, in a cluster of *v* distinct molecules. As with the single-chain case, the GCT introduces a few key variables. These are defined in terms of 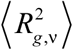 the ensemble-averaged mean-squared radius of gyration of a single chain within a cluster of *v* molecules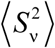 the ensemble-averaged mean-squared radius of gyration of a cluster of *v* molecules, and α_s,*v*_ the contraction ratio of single chain within the cluster. Accordingly,

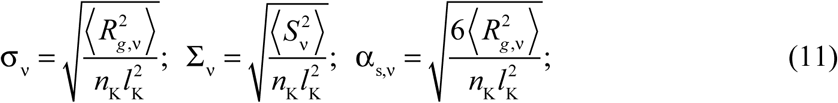

For a given value of τ, the free energy *A*_*v*_ of a cluster of *v* molecules, with 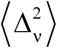 as the ensemble-averaged mean-squared distance between chains in the cluster, and *q* as the value of the compression parameter for a single chain, is the sum of three terms:

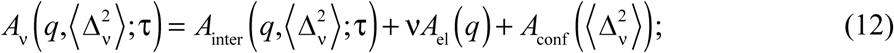

The contact free energy *A*_inter,*v*_ combines contributions from two- and three-body contacts among chain segments. Using the assumption of a Gaussian distribution for *W*(*R*_*i*_) it follows that:

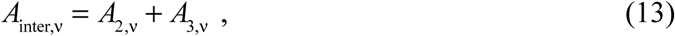

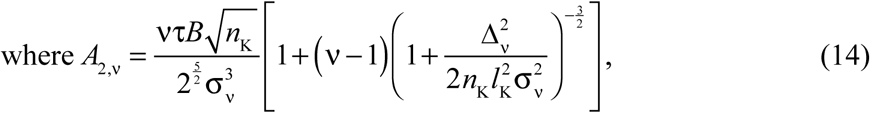

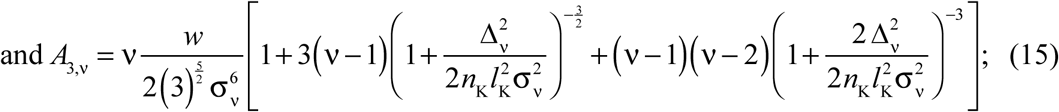

*vA*_el_ is the elastic free energy of individual chains in the cluster whereas *A*_conf_ is the free energy cost associated with confining *v* molecules into a cluster. The confinement term accounts for the loss of translational entropy. It also includes a combinatorial Gibbs-like correction term that accounts for the indistinguishability of chains within a cluster. The expression for *A*_conf_ derived by Raos and Allegra, is as follows:

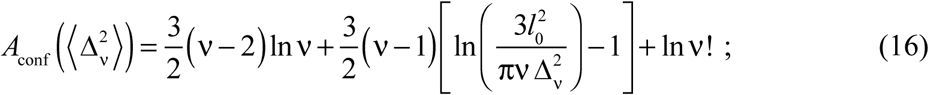

Phase separation is an infinitely cooperative process [66] and accordingly, the cluster free energy is minimized in the limit *v* → ∞. In this limit, it is useful to work in terms of the volume fraction ϕ. Raos and Allegra show that the free energy per chain in a polymer solution *a*_ch_ is written in the limit *v* → ∞ as:

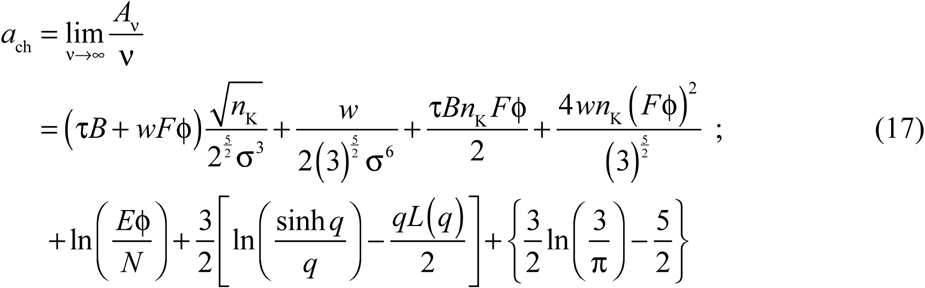

Here, 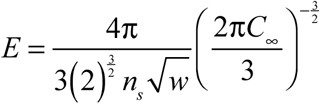 and 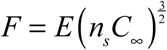. The expression in equation (17) is then used to estimate the free energy per unit volume in the polymer solution denoted as *a*_vol_. Here, 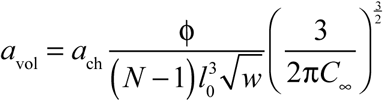.

For specific values of ϕ and τ there should be a single value of *q* that minimizes the system free energy. This is estimated from a generalization of the expression in equation (9) now rewritten for a single chain in a polymer solution as:

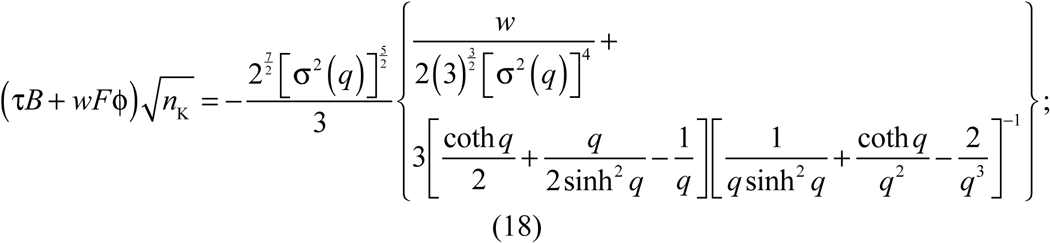

In order to compute the phase diagram, we need an expression for the chemical potential µ_P_ of the polymer as a function of values of ϕ. The chemical potential is the partial derivative of *a*_vol_ with respect to the number of polymer molecules. This takes the form:

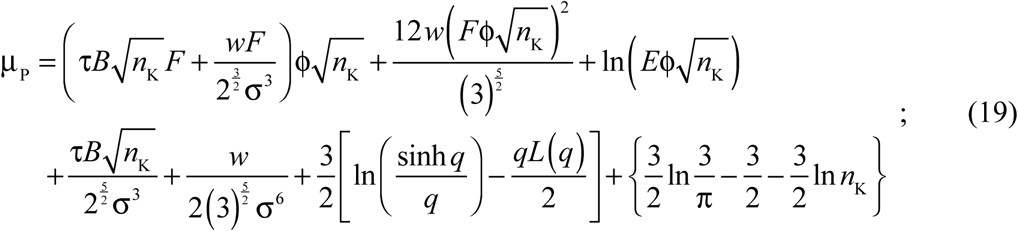

For fixed values of τ, *B*, and *w*, the chemical potential is a function of the compression parameter *q* and the volume fraction ϕ. Since there is a one-to-one numerical mapping between *q* and ϕ, it follows that we can calculate µ_P_ for different values of ϕ given prior knowledge of the two- and three-body interaction coefficients *B* and *w*, respectively.

### Calculation of full phase diagrams

The mapping between *q* and ϕ is extracted numerically from equation (17). For this mapping we use values of *B* and *w* derived from the analysis of single-chain coil-to-globule transitions. Next, for fixed τ, we calculate the locations of spinodals by computing the values of ϕ that are solutions of the equation 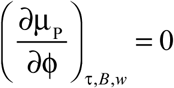. This yields two solutions *viz*., ϕ_s1_(τ) and ϕ_s2_(τ). Next, to generate the coexistence curve, we compute the locations of coexisting dilute and dense phases *viz*., ϕ_b1_(τ) and ϕ_b2_(τ). For fixed τ, the two coexisting phases represent points for which the chemical potential (µ_P_) and osmotic (∏) pressure are equalized along the horizontal tie line that connects the points ϕ_b1_ and ϕ_b2_. Accordingly, it follows that:

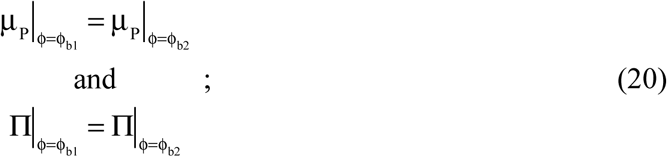

The osmotic pressure is calculated using the expression:

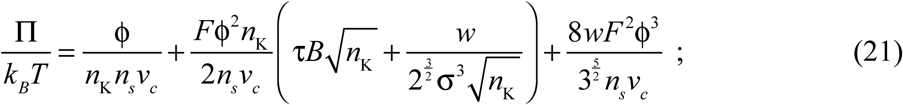

Here, 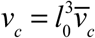 and 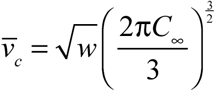. For fixed τ, *B*, and *w*, we use the expressions in equations (19) – (21) and numerically identify the points ϕ_b1_ and ϕ_b2_ that lie on the same tie line. The locus of points ϕ_b1_(τ) and ϕ_b2_(τ) obtained for different values of τ yield the desired binodal in the (ϕ,τ) or (ϕ,*T*) plane.

## RESULTS

We demonstrate the application of the two-pronged approach wherein we obtain system-specific parameters from coarse-grained or all-atom simulations of coil-to-globule transitions of individual and incorporate them into the GCT to obtain full phase diagrams. Here, full phase diagrams refer to system-specific binodals and spinodals that are calculated and shown in the two-parameter space with volume fraction ϕ along the abscissa and simulation temperature *T* along the ordinate. Note that *T* is unitless for phase diagrams computed using PIMMS since it is written in terms of *k*_*B*_*T*, which was set to one by Martin et al. [10]. For phase diagrams computed using all-atom simulations, the temperature is in units of degree-Kelvin.

### Phase diagrams for Aro^+^, WT-A1, Aro^-^, and Aro^--^ variants

First, we extract the theta temperature by analyzing the internal scaling profiles obtained from PIMMS simulations of coil-to-globule transitions for each of the four systems. This analysis is summarized in **Figure S2**. Additionally, **Figure S3** shows plots of 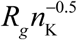 as a function of τ for each of the four constructs. The system-specific estimates for *T*_θ_ are summarized in each of the panels of **Figure 4**. Each panel in **Figure 4** also shows a system-specific profile for the contraction ratio plotted as a function of 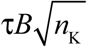. **Figure 5** shows a comparison between binodals obtained using GCT and those obtained directly from PIMMS simulations. The latter were shown by Martin et al. [10] to reproduce relative trends measured using experiments and can be brought into the same currency as the experiments using a multiplicative factor for the temperature and a conversion from volume fractions into molar units. The results shown in **Figure 5** indicate that the binodals obtained using GCT, based on the parameters extracted from single-chain simulations, are in good agreement with binodals obtained from direct simulation of phase separation using PIMMS. The critical temperature as well as the width of the two-phase regimes decrease with decreasing sticker valence and these trends are accurately captured by the GCT derived binodals.

**Figure 4:**
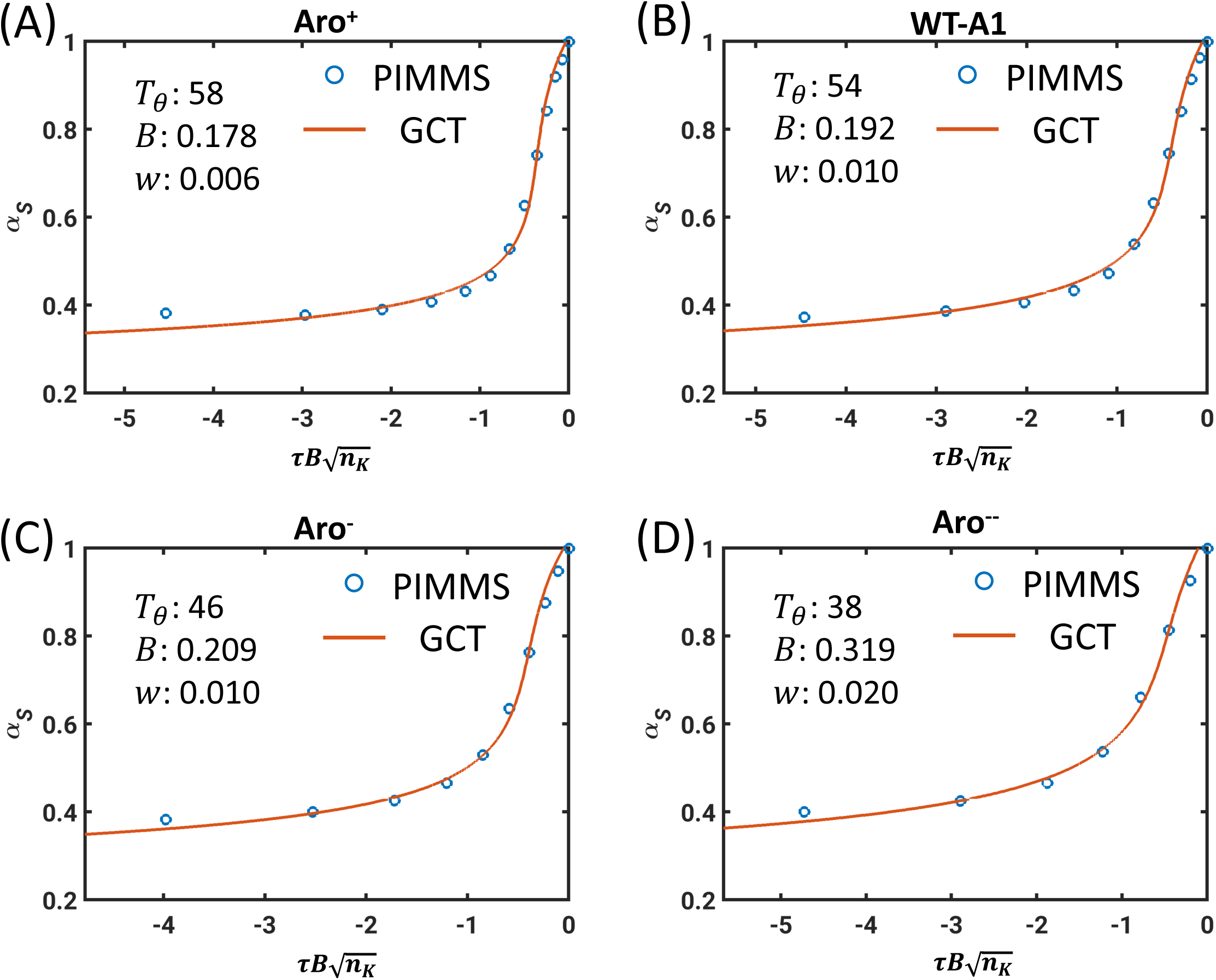
Profiles of contraction ratios for Aro^+^, WT-A1, Aro^-^, and Aro^--^. In each panel, the blue circles are results from PIMMS simulations of single-chains and the red curves are contraction ratios obtained using equation (9) that best fits the simulation results. The best fit, system-specific parameters for *T*_θ_, *B*, and *w* are shown as legends in each of the four panels. *B* has units of inverse volume and *w* has units of the square of the inverse volume.

**Figure 5:**
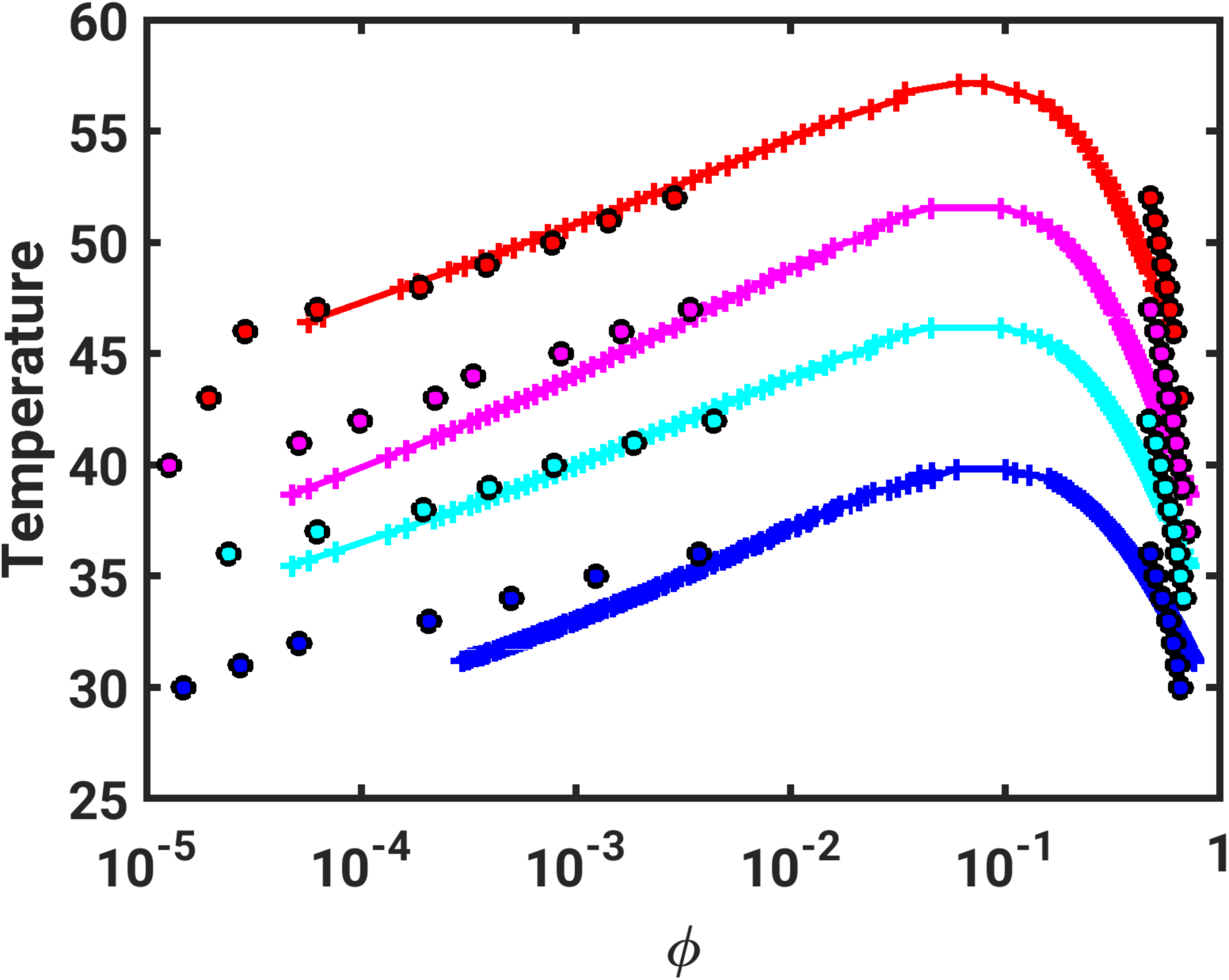
Comparison of binodals obtained using direct simulations based on PIMMS and numerical implementation of the GCT formalism that uses parameters extracted from single-chain simulations. The PIMMS results are shown as circles and the results using GCT are shown as + marks. Here, the binodals for the different systems are shown using symbols of different colors red (Aro^+^), magenta (WT-A1), cyan (Aro^-^), and blue (Aro^--^). The binodals obtained using GCT were shifted upward along the ordinate by 10.5 units and along the abscissa by –0.3 units to bring them onto the same scale as the PIMMS derived binodals.

Direct simulations of phase separation enable the extraction of sequence- or architecture-specific phase diagrams [10, 16, 20, 26]. However, identifying the critical point is challenging because the amplitudes of conformational fluctuations become commensurate with chain dimensions as *T* approaches *T*_c_. Similarly, as ϕ approaches ϕ_c_, the amplitudes of concentration fluctuations become equivalent or even larger than the simulation volume. It is also challenging to estimate the low and high concentration arms of spinodals as this requires direct observation of the onset of instabilities. These challenges are readily overcome in the GCT-aided calculation of phase diagrams.

Details regarding the width of the metastable region and its dependence on *T* are direct determinants of the mechanisms of phase separation [14]. For a given *T* < *T*_c_, if ϕ_0_(*T*) is the initial value of ϕ and ϕ_b1_(*T*) < ϕ_0_(*T*) < ϕ_s1_(*T*), then the system is in the metastable regime. The rate of nucleation of phase separation from an initial homogeneous one-phase state will be governed by two parameters *viz*., 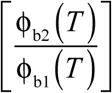, which is a measure of the width of the two-phase regime, and 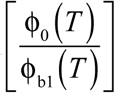, which is a measure of the supersaturation [67]. An analogous relationship exists for the initial conditions ϕ_s2_(*T*) < ϕ_0_(*T*) < ϕ_b2_(*T*), in which the starting conditions are in the high-concentration metastable regime. For simplicity we will focus here on the low-concentration metastable regime, since this is the most relevant scenario for biological systems. If ϕ_s1_(*T*) < ϕ_0_(*T*) < ϕ_s2_(*T*), then the dynamics of phase separation follow spinodal decomposition [14, 68]. These dynamics are best modeled using the Cahn-Hilliard formalism [69], although a convolution with sol-gel transitions that dynamically arrests phase separation cannot be ignored [32, 70]. The quench depth relative to the binodal and spinodal directly determines the dynamics of phase separation. Knowledge of the quench depth helps one discern whether the dynamics can be modeled using a Cahn-Hilliard formalism [71, 72].

Calculations of full phase diagrams should aid in the assessment of comparative dynamics for similar quench depths across different systems and for different quench depths of the same system. Panel (A) in **Figure 6** shows the GCT-derived binodals and spinodals for WT-A1. The predicted spinodal is manifest only for larger volume fractions *viz*., ϕ > 10^−2^, which would correspond to initial concentrations that are in the millimolar range for WT-A1. Accordingly, concentrations corresponding to initial conditions in typical *in vitro* experiments are likely to lie between the binodal and spinodal thereby implicating nucleation as the dominant mechanism for phase separation. Further, the low endogeneous concentrations that one expects for *in vivo* settings are also likely to correspond to the metastable regime.

**Figure 6:**
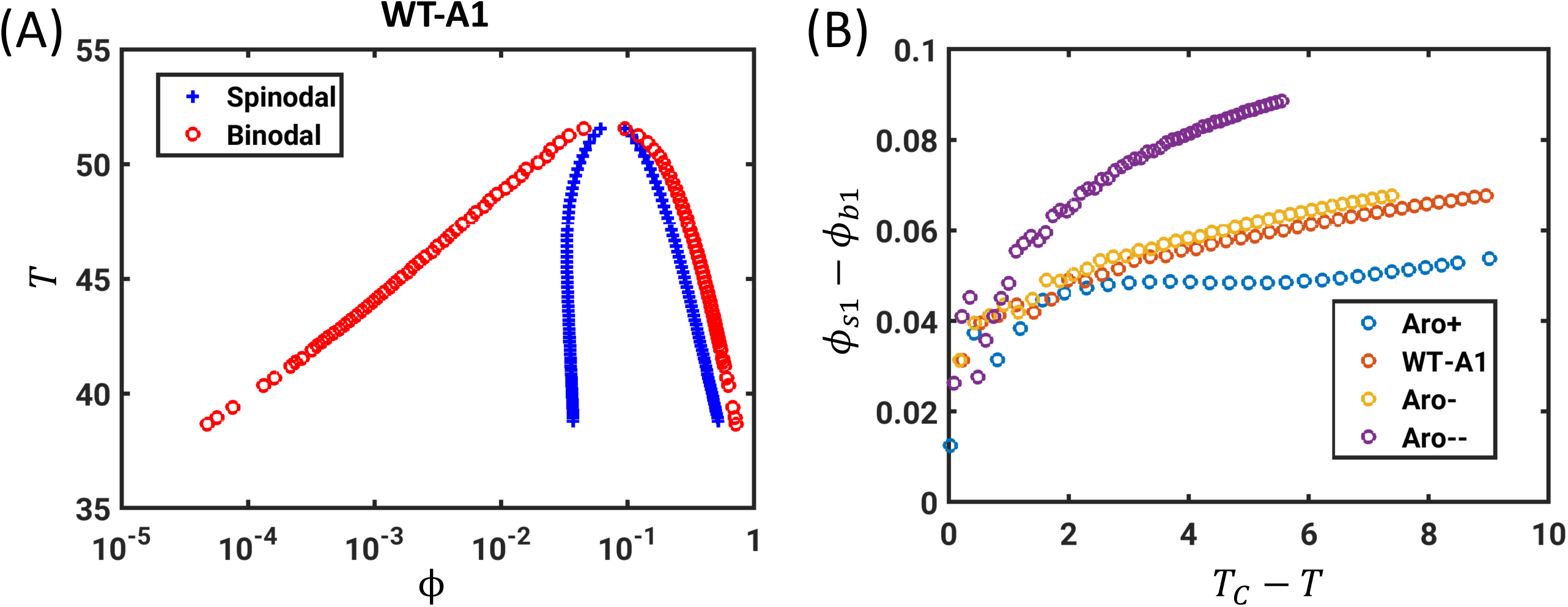
Assessments of the binodals vs. spinodals for the WT-A1 and three designed variants. (A) Full phase diagram, binodal and spinodal, for the WT-A1 system. Here, we use the unitless simulation temperatures from PIMMS. (B) Analysis of how the width of the metastable regime Δϕ_m_(*T*) varies with (*T – T*_c_) for each of the four sequences.

Panel (B) in **Figure 6** quantifies how the width of the metastable region varies with temperature. Specifically, this plot shows how Δϕ_m_ = [ϕ_s1_(*T*) – ϕ_b1_(*T*)] varies with (*T*_c_ – *T*) for Aro^+^, WT-A1, Aro^-^, and Aro^--^, respectively. As expected, the width of the metastable region Δϕ_m_ approaches zero as *T* approaches *T*_c_. In general, for (*T*_c_ – *T*) > 2, Δϕ_m_ is on the order of 10^−2^ and for a given value of *T*, Δϕ_m_ is smallest for Aro^+^ and largest for Aro^--^. Additionally, Δϕ_m_ varies most steeply with (*T*_c_ – *T*) for Aro^--^, followed by Aro^-^, WT-A1, and Aro^+^, respectively. These results indicate that the width of the metastability region decreases and shows a weaker dependence on temperature as the system becomes a stronger driver of phase separation.

### Phase diagrams for polyQ and synthetic IDPs

The results shown in **Figure 5** illustrate the accuracy of the GCT derived binodals as compared to binodals obtained using PIMMS for four closely related systems. Next, we computed full phase diagrams using parameters extracted from all-atom simulations that were based on the ABSINTH implicit solvation model. We present results for three separate systems *viz*., polyQ and two synthetic IDPs, one with UCST behavior and another showing LCST behavior. The results for the polyQ system are shown in **Figure 7**.

**Figure 7:**
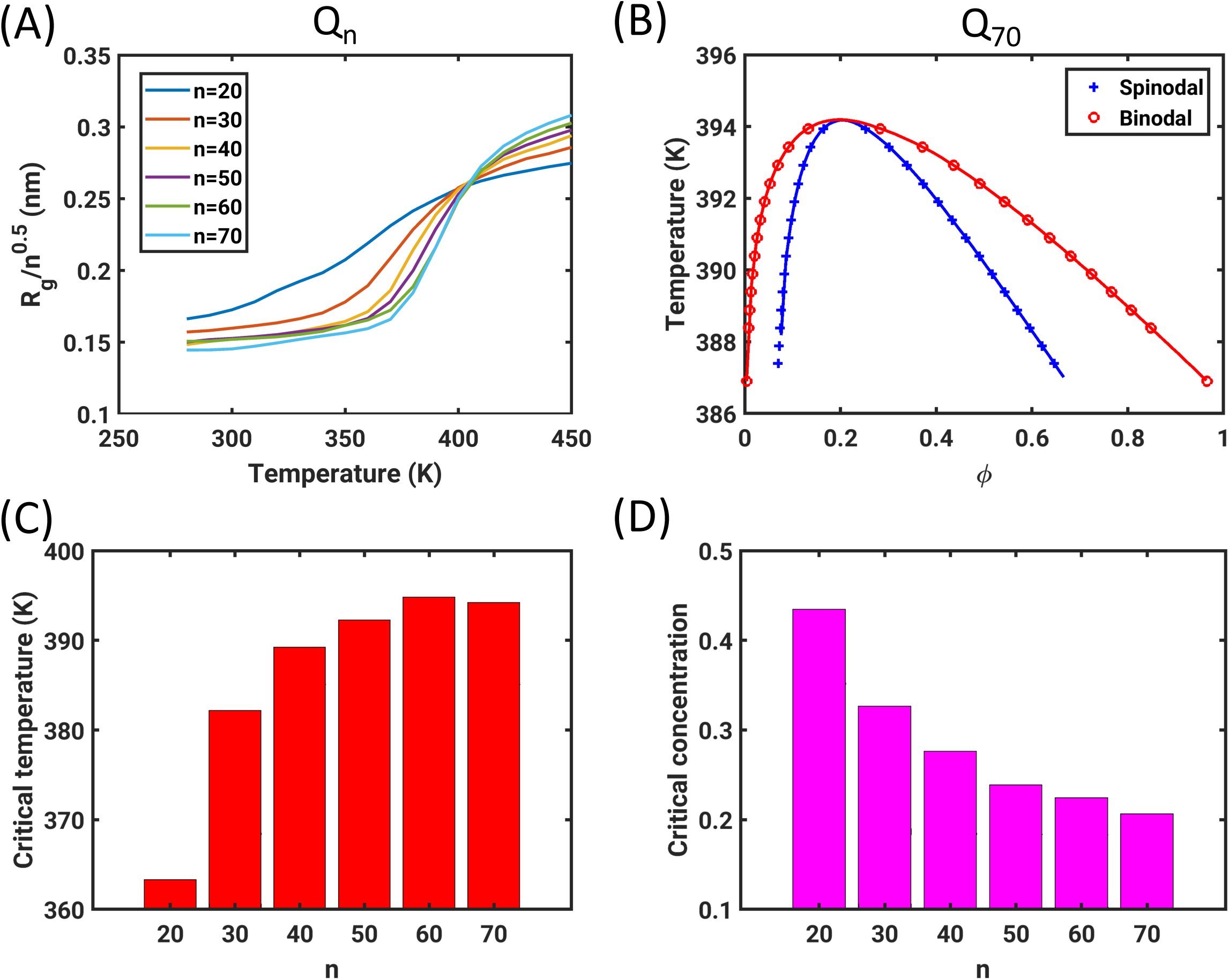
Results for the polyQ system. (A) Normalized *R*_g_ for different chain lengths as a function of simulation temperature *T*. The theta temperature (*T*_θ_ ≈ 405K) corresponds to the point where the curves coincide with one another. Note that previous simulations based on the ABSINTH model estimated a value of 390K for the *T*_θ_ of polyQ [41, 57]. The discrepancy between published and current estimates come from our current usage of a temperature dependent functional form for the macroscopic dielectric constant of water [56]. (B) Full phase diagrams for the Q_70_ system. The binodal and spinodal coincide at the critical point. The smooth red and blue curves join the dots along the binodal and spinodal, respectively and they are used to guide the eye. (C) Bar plot showing the increase of the critical temperature (*T*_c_) with polyQ length. (D) Bar plot showing the decrease of the critical volume fraction (ϕ_c_) with increasing polyQ length.

Crick et al. previously measured the temperature dependence of the solubility limit for different constructs with polyQ tracts [34]. The measured saturation concentrations, which are in the low micromolar range, decrease to be in the sub-micromolar range for longer polyQ molecules. These measured concentrations translate to volume fractions that are ∼10^−5^ and are what we obtain for the low arms of binodals (**Figure 7**). Further, Crick et al. showed that polyQ has UCST behavior with a critical temperature that is above the boiling point of water – a feature that is recapitulated by the simulations and the GCT-aided calculations. Simulation results, summarized in panel (A) of **Figure 7**, were used to identify the theta temperature for this system. Here, *T*_θ_ ≈ 405K. Panel (B) shows the full phase diagrams, including binodals and spinodals for phase separation. These data are shown for the Q_70_ system. Panels (C) and (D) show that the critical temperature (*T*_c_) increases with increasing polyQ length whereas the critical volume fraction (ϕ_c_) decreases with increasing polyQ length. This is in accord with general trends expected from mean-field theories for the phase separation of homopolymers [14].

Finally, **Figure 8** summarizes results obtained for the two synthetic IDPs with UCST and LCST behavior. Recent attention has focused on IDPs derived from resilin- and elastin-like polypeptides that are designed to have UCST versus LCST behavior, respectively [29, 73]. We show results for two archetypal systems. The theta temperatures for these systems were estimated by analyzing the temperature dependent internal scaling profiles. Panels (A), (B), and (C), respectively show the profile of the normalized *R*_g_ plotted as a function of simulation temperature, the contraction ratio as a function of 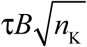, and the resultant phase diagram for the (QGQSPYG)_9_ system that shows UCST behavior. Panels (D), (E), and (F) show the corresponding results for the (TPKAMAP)_9_ system that has LCST phase behavior. Systems that are perfect repeats of specific peptide sequences may be thought of as homopolymers on length scales that are longer than one repeat. Accordingly, these systems are amenable to a joint computational approach that combines simulations of single chains with GCT to extract computed phase diagrams. The results demonstrate that the GCT is applicable for calculating phase diagrams of systems with LCST behavior. All that is required is that simulations at the single chain level demonstrate a collapse transition at higher temperatures, which we observe for the (TPKAMAP)_9_ system if we use temperature-dependent reference free energies of solvation as prescribed by Wuttke et al. [56]– see panel (D) of **Figure 8**.

**Figure 8:**
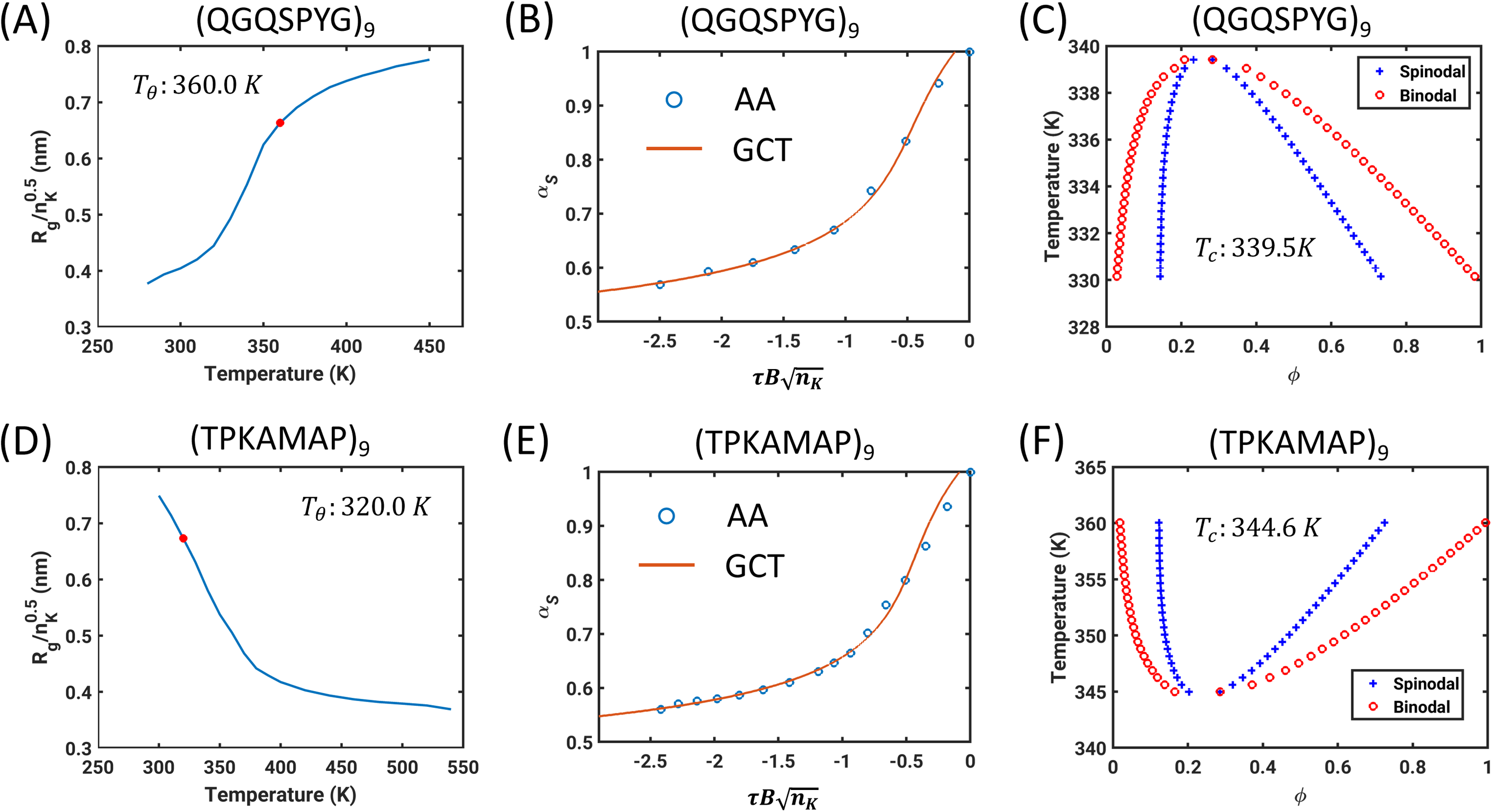
Results for synthetic systems that show UCST behavior (top row) and LCST behavior (bottom row). (A) Normalized *R*_g_ plotted as a function of simulation temperature for (QGQSPYG)_9_. The intercept along the abscissa from the red circle corresponds to the location of the theta temperature. (B) Contraction ratio plotted against 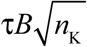 for (QGQSPYG)_9_. (C) Binodal and spinodal for (QGQSPYG)_9_. The critical point is where the binodal and spinodal coincide and the estimated UCST is *T*_c_ ≈ 340 K. The symbols represent the temperatures for which the points on the binodal and spinodal were obtained from the GCT theory. (D) Plot that is equivalent to (A) for the system (TPKAMAP)_9_ that has LCST behavior. This system undergoes a robust collapse transition above the theta temperature, which is the intercept along the abscissa that passes through the red circle. (E) Plot of the contraction ratio for (TPKAMAP)_9_ as a function of 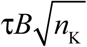. Note that *B* is positive for UCST systems and negative for LCST systems. Accordingly, 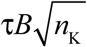 is always negative. (F) Full phase diagram, including the binodal and spinodal for (TPKAMAP)_9_. The LCST (≈ 345 K) corresponds to the temperature where the binodal and spinodal coincide.

## DISCUSSION

In this work we have adapted and deployed the Gaussian cluster theory of Raos and Allegra [12, 13] in order to compute full phase diagrams for archetypal LCDs using parameters extracted from single-chain simulations of coil-to-globule transitions. Our motivation for adapting and deploying the GCT-aided approach comes from recent observations showing that mean-field theories based on the Flory-Huggins theory [74, 75] can be reliably used to describe experimentally measured binodals for LCDs [3, 8-10]. Bioinformatics analyses have identified a uniform linear distribution of stickers as a defining feature of LCDs that are implicated in liquid-liquid phase separation [10, 76, 77]. This suggests that, to zeroth order, LCDs that are drivers of liquid-liquid phase separation can be treated as effective homopolymers. Additionally, recent theoretical, computational, and experimental work has shown that there is a one-to-one correspondence between the driving forces for chain collapse in dilute solutions and phase separation in concentrated solutions of LCDs [10, 16, 26, 65, 78, 79]. The theoretical work of Lin and Chan, based on a random phase approximation based mean field approach to incorporate the effects of sequence patterning and cation-pi as well as pi-pi interactions, showed an inverse correlation between the extent of collapse in polyampholytes and the critical temperature [78]. Overall, the observations to date are suggestive of the swapping of intra-chain interactions for inter-chain interactions as being a key determinant of phase separation [15]. Therefore, we reasoned that a theory designed to connect single chain coil-to-globule transitions to phase separation should be suitable for numerical adaptation. We selected the GCT for our two-pronged approach because, to the best of our knowledge, this is one of the few theories that describes both single-chain conformational equilibria and multi-chain phase equilibria using a single set of parameters.

Use of the GCT approach requires *a priori* numerical characterization of single chain coil-to-globule transition profiles. These can be extracted from all-atom or coarse-grained simulations, which are accessible via different approaches. Computational methods based on the ABSINTH implicit solvation model and novel refinements to explicit solvent models have enabled increasingly accurate and efficient simulations of conformational equilibria of a range of IDPs [36, 37, 56, 60, 80-96]. Many of these approaches can use experimental data as inputs to generate accurate coarse-grained descriptions for conformational ensembles of IDPs [97, 98]. These methods can be brought to bear for accurate characterization of coil-to-globule transitions. Here, we focused on temperature as the control parameter; however, one can conceive of other intensive parameters such as pH [99] or the chemical potentials of solution ions or small molecules as control parameters [8, 11, 18, 100].

Our method of extracting parameters from single-chain coil-to-globule transitions and computing phase diagrams using a numerical approach based on the GCT is similar in spirit to the field-theoretic approaches that have been adapted from the polymer literature for the study of LCD phase separation [101, 102]. The essential differences are that the GCT, unlike field-theoretic approaches, remains primarily a mean-field theory. This implies that effects of fluctuations are ignored in the GCT formalism. This is likely to create issues in describing how the width of the two-phase regime varies as the critical temperature is approached [103]. However, as a rapid and potentially easy to deploy strategy, the GCT should afford a useful approach for high-throughput comparative assessments of phase diagrams providing we have results from simulations of coil-to-globule transitions. This GCT-driven approach is likely to be useful for the design of repetitive sequences with bespoke phase behavior – a problem of considerable interest in the design of novel biomaterials as well as synthetic condensates. It can be combined with field-theoretic approaches to obtain improved descriptions of near critical regions and of the effects of density fluctuations engendered by conformational fluctuations.

Our use of the GCT rests on the validity of the assumption that the interactions that drive single chain collapse in dilute solutions are the same as the ones that drive inter-chain associations in concentrated solutions. We refer to this as the *strong coupling regime* and note that the correspondence required for the use of GCT need not always prevail [18, 103]. For example, a direct prediction of the GCT, which we have not called out here explicitly, is the fact that the chain conformations in the dense phase are likely to be distinct from those in the coexisting dilute phase. Well below the theta temperature, chains in dilute solutions should form globules, whereas chains in the dense phase are likely to behave like random walks because intermolecular interactions weaken the driving force for collapse [13]. Therefore, systems characterized by limited conformational change across the phase boundary are poor candidates for the use of GCT. This will be true as we approach the critical temperature, where the two-phase regime narrows giving rise to ultra-dilute droplet phases, which has been observed in recent studies [18].

Predictions from the GCT can be tested by explicit measurements of binodals and by characterizing the dimensions of individual chains in both the dilute and dense phases as a function of temperature or any other control parameter of interest. This should be feasible using advances in small angle x-ray scattering as well as multi-parameter single molecule fluorescence spectroscopies [91, 104, 105].

## SUPPORTING MATERIAL

The supporting information appendix includes Figures S1 – S3, Tables S1 and S2, and documented MATLAB code (https://www.mathworks.com) for extracting *T*_θ_, *B*, and *w* from analysis of temperature-dependent profiles for contraction ratios for each of the examples presented here.

## AUTHOR CONTRIBUTIONS

Conceptualization: XZ and RVP; All-atom simulations and GCT implementation: XZ; PIMMS simulations: ASH; Analysis: XZ and R.V.P; Writing: RVP; Editing: XZ, ASH, AC, TM, and RVP; Funding: AC, TM, and RVP.

## ACKNOWLEDGMENTS

This work was supported by grants DMR 1729783 from the US National Science Foundation (to A.C. and R.V.P) and the St. Jude Children’s Research Collaborative on Membraneless Organelles (to T.M. and R.V.P). We thank Mina Farag for helpful discussions and critical inputs.

